## Supplementary material for "Mechanically stimulated osteocytes maintain tumor dormancy in bone metastasis of non-small cell lung cancer by releasing small extracellular vesicles"

### Supplemental Figures

#### Supplemental Figure 1

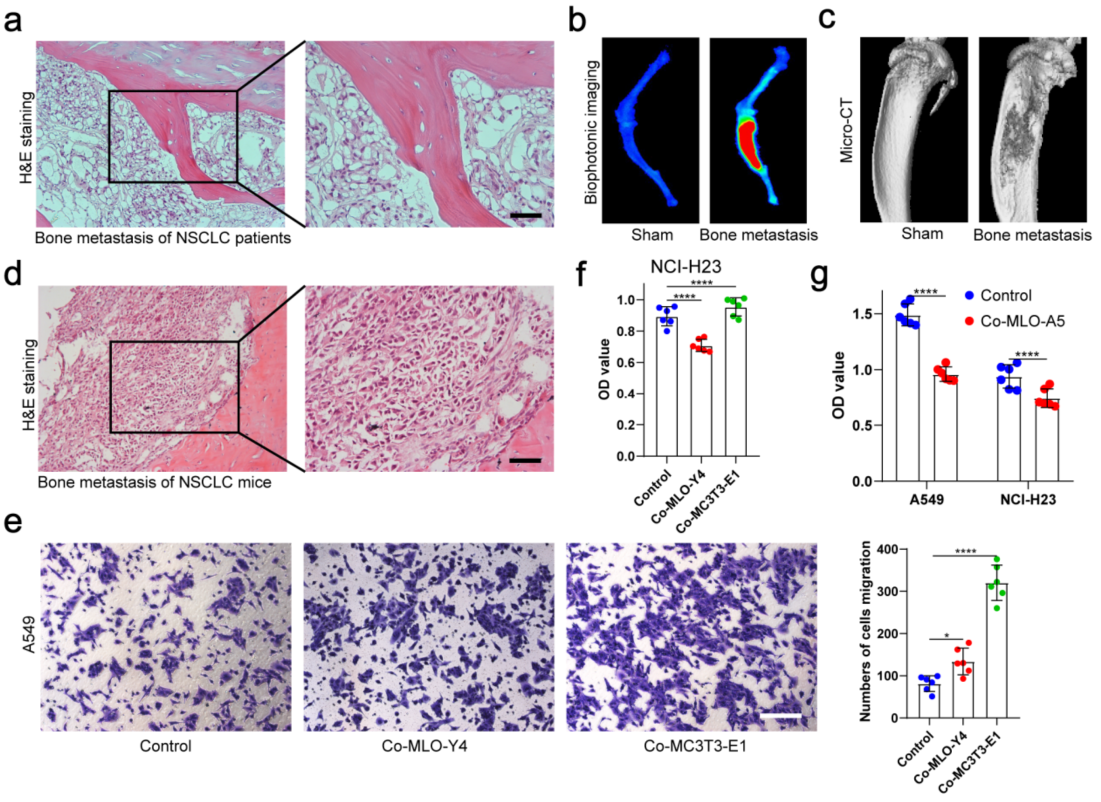


**Supplemental Figure 1. Histological identification of patients and mice with bone metastases and effects of MLO-Y4 and MC3T3-E1 on proliferation and migration of NSCLC cells.**

a. The representative images of H&E staining in bone metastatic sites of patients with NSCLC. Scale bars: 50 μm.

b. An intraosseous model of bone metastasis was used via direct implantation of A549 cells expressing green fluorescent protein (GFP) into tibia of mouse. Representative biophotonic images of NSCLC cells in hindlimb bones of mice.

c. Representative micro-CT imaging of the tibia in mice with or without bone metastasis.

d. The representative images of H&E staining of bone metastasis in tibia of mice. Scale bars: 50 μm.

e. Transwell assays were performed to evaluate the effects of MLO-Y4 cells or MC3T3-E1 cells on the migration of A549 cells in the co-culture system. Scale bars: 100 μm. n=6, one-way analysis of variance with Turkey's multiple comparisons test.

f. CCK-8 assays were performed to evaluate the effects of MLO-Y4 cells or MC3T3-E1 cells on the proliferation of NCI-H23 cells in the co-culture system. n=6, one-way analysis of variance with Turkey's multiple comparisons test.

g. CCK-8 assays were performed to evaluate the effects of MLO-A5 cells on the proliferation of A549 and NCI-H23 cells in the co-culture system. n=6, student's two-sided unpaired t test.

no significance(ns) $P>0.05$, *$P < 0.05$, **$P < 0.01$, ***$P < 0.001$, ****$P < 0.0001$.

**Supplemental Figure 2**

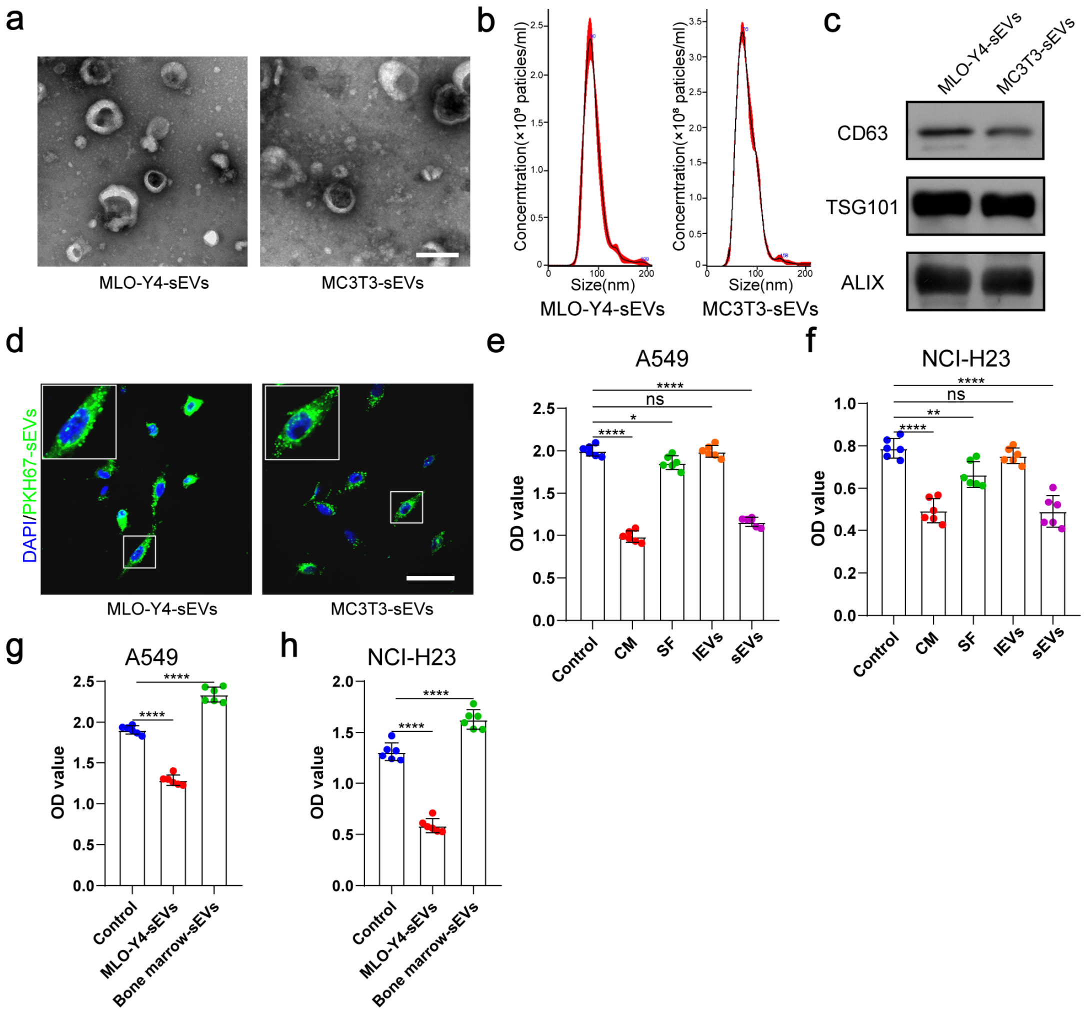


**Supplemental Figure 2. Identification of MLO-Y4-sEVs and MC3T3-E1-sEVs and their effects on the proliferation ability of NSCLC cells.**

Conditioned medium (CM) was separated into soluble factor (SF), large extracellular vesicles (lEVs), and small extracellular vesicles (sEVs) fractions, by serial ultracentrifugation and ultrafiltration.

a. Representative TEM images of sEVs extracted from conditioned medium. Scale bars, 100 nm.

b. NanoSight particle analysis displays the size distribution of the sEVs.

c. Western blot analysis of the typical sEVs markers (CD9, Alix, and TSG101) of the sEVs isolated from the culture medium.

d. A549 cells were incubated with PKH67-labeled sEVs and subjected to immunofluorescence of PKH-67-labeled sEVs (green) and DAPI (blue) for nuclei. Scale bars, 50 μm.

e, f. CCK-8 assays were performed to evaluate the effects of the components of the MLO-Y4 cells culture medium on the proliferation of NSCLC cells. n=6, one-way analysis of variance with Turkey's multiple comparisons test.

g, h. CCK-8 assays were performed to evaluate the effect of MLO-Y4-sEVs or bone marrow-sEVs on the proliferation of NSCLC cells. n=6, one-way analysis of variance with Turkey's multiple comparisons test.

no significance(ns) $P>0.05$, *$P < 0.05$, **$P < 0.01$, ***$P < 0.001$, ****$P < 0.0001$.

**Supplemental Figure 3**

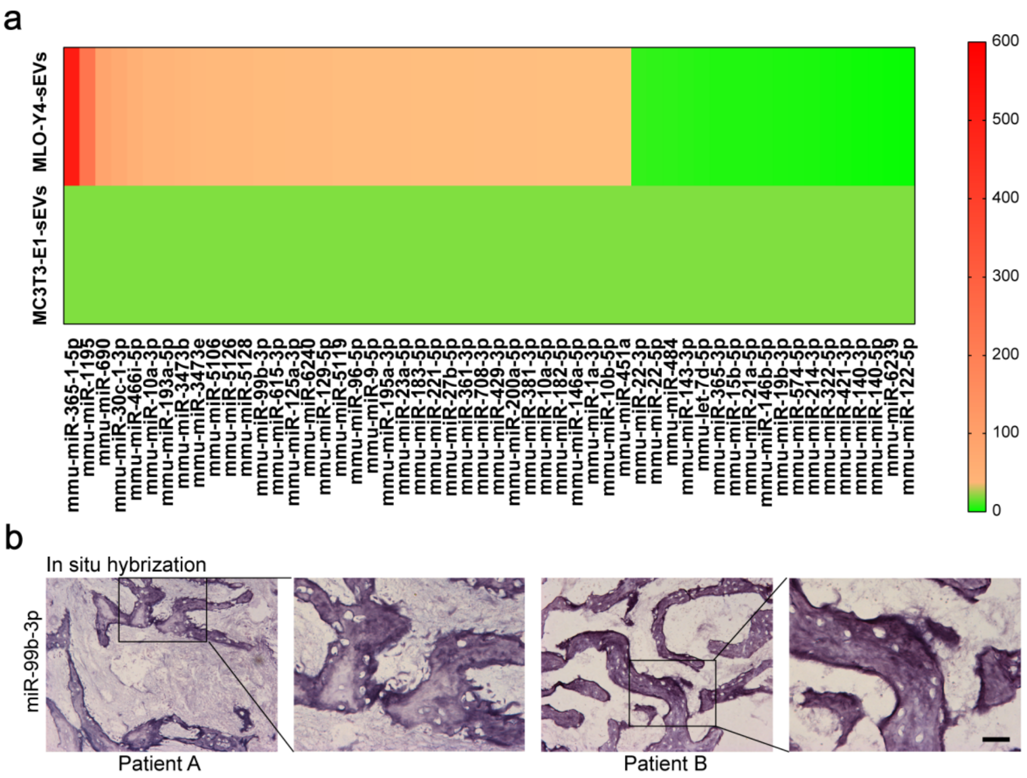


**Supplemental Figure 3. The miRNA profiles of MLO-Y4-sEVs and MC3T3-E1-sEVs and distribution of miR-99b-3p in bone metastases tissues of NSCLC patients.**

a. The heat map shows differentially expressed miRNAs in sEVs at least 1.5-fold difference between MLO-Y4-sEVs and MC3T3-E1-sEVs identified by microarray.

b. Representative in situ hybridization images showing the expression of miR-99b-3p in bone metastatic sites of patients with NSCLC. Scale bars, 50 μm.

#### Supplemental Figure 4

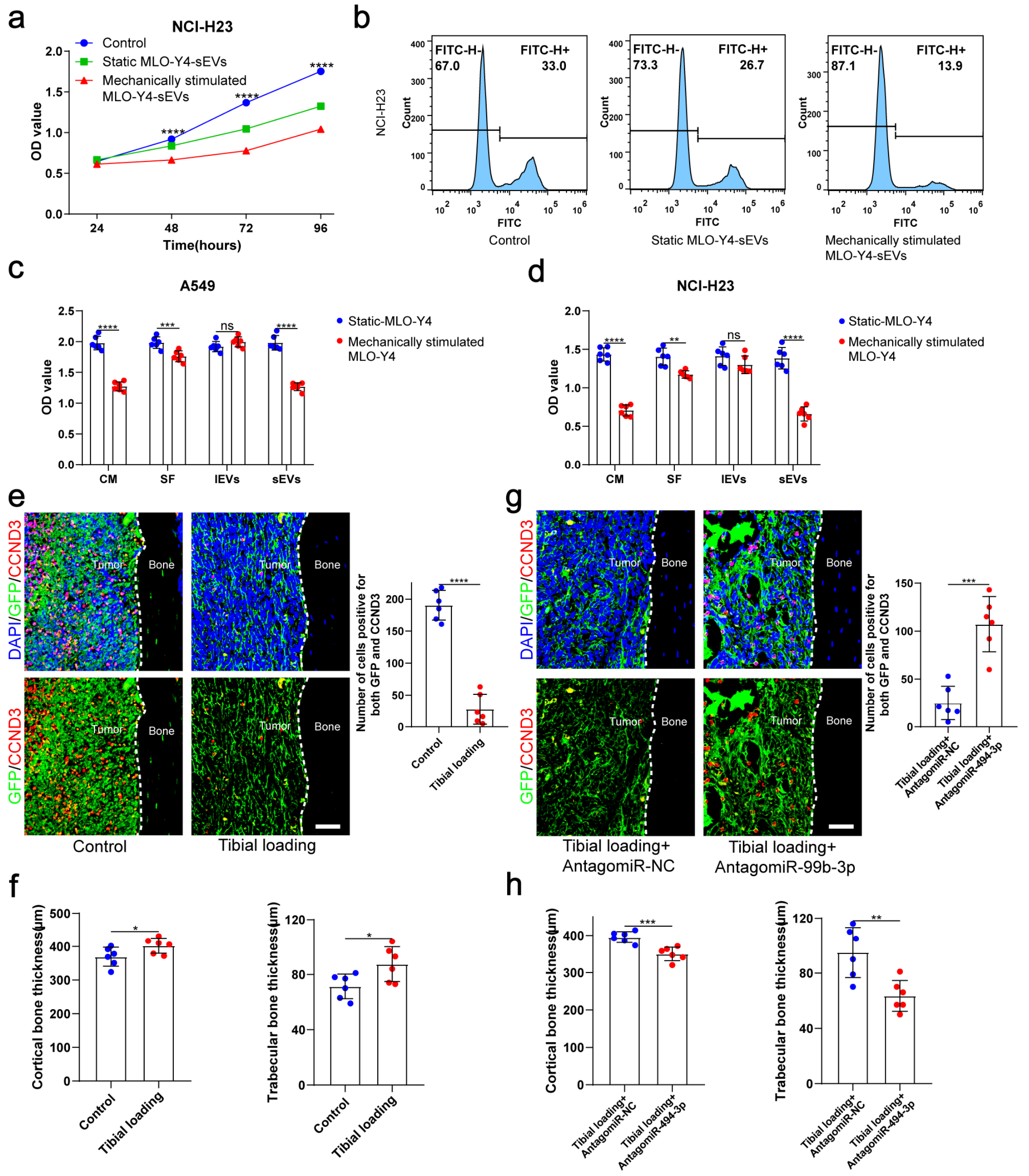


**Supplemental Figure 4. Mechanical loading increased the release of sEVs from osteocytes and enhanced inhibitory effect of osteocytes on the proliferation of NSCLC cells.**

a. MLO-Y4 cells were mechanically stimulated by stretching at 12% strain at a 1.25 Hz frequency for 12 h, and sEVs were extracted from the conditioned medium. CCK-8 assays were performed to evaluate the effect of sEVs on the proliferation of NCI-H23 cells. n=4, two-way analysis of variance with multiple comparisons test.

b. EdU flow cytometry were performed to evaluate the effect of sEVs on the proliferation of NCI-H23 cells.

c, d. CCK-8 assays were performed to evaluate the effects of the components of the static or mechanically stimulated MLO-Y4 cells culture medium on the proliferation of NSCLC cells. n=6; one-way analysis of variance with Turkey's multiple comparisons test.

e, g. Representative immunofluorescence staining images of GFP (green), CCND3 (red), and DAPI (blue) in the tibia of mice with bone metastases and quantification of the GFP and CCND3 positive cells. The dashed line indicates the boundary between bone and tumor. Scale bars: 50 μm. n=6, one-way analysis of variance with Turkey's multiple comparisons test.

f, h. Representative micro-CT imaging and quantitation of the tibia (cortical bone thickness, trabecular bone thickness) in mice. n=6, one-way analysis of variance with Turkey's multiple comparisons test.

no significance(ns) P>0.05, *P < 0.05, **P < 0.01, ***P < 0.001, ****P < 0.0001.

**Supplemental Figure 5**

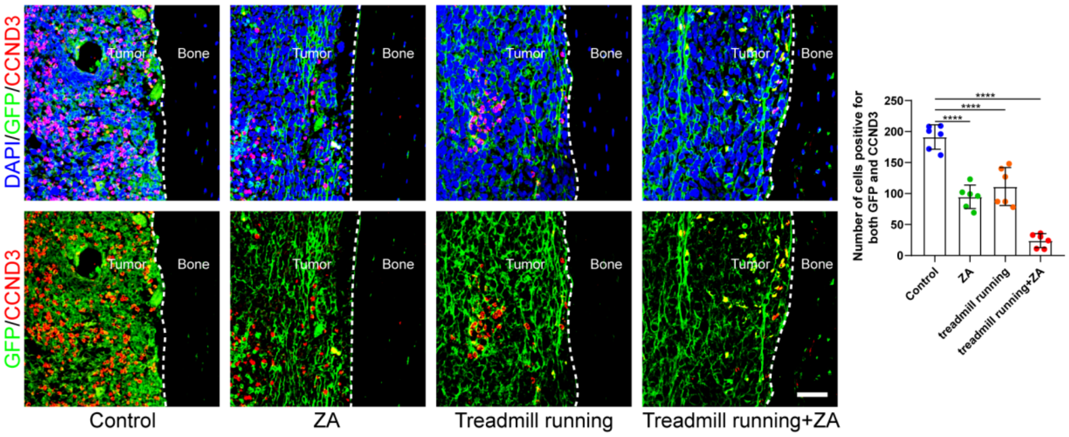


**Supplemental Figure 5. Moderate exercise combined with zoledronic acid effectively suppressed the progression of bone metastasis of NSCLC.**

Representative immunofluorescence staining images of GFP (green), CCND3 (red), and DAPI (blue) in the tibia of mice with bone metastases and quantification of the GFP and CCND3 positive cells. The dashed line indicates the boundary between bone and tumor. Scale bars: 50 µm. n=6, one-way analysis of variance with Turkey's multiple comparisons test.

**Supplemental Figure 6**

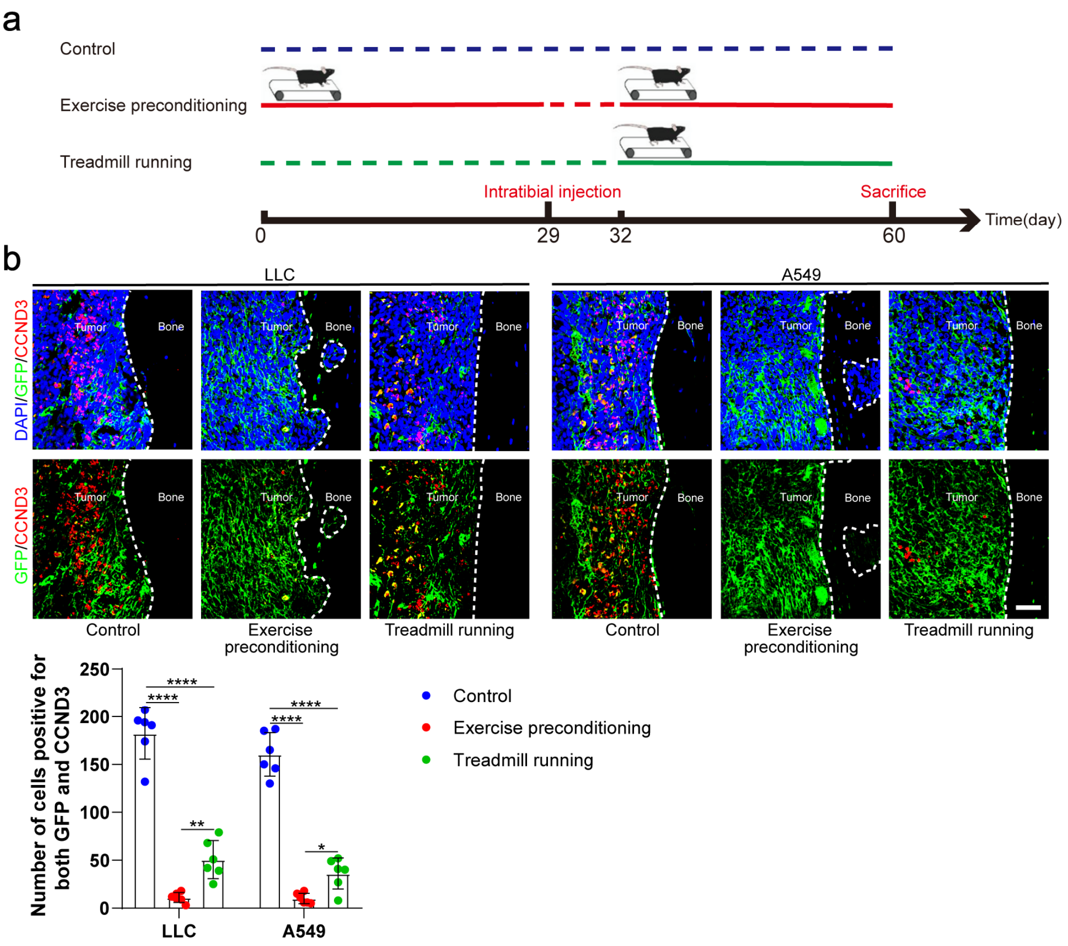


**Supplemental Figure 6. Exercise preconditioning effectively suppressed the progression of bone metastasis of NSCLC.**

a. An intraosseous model of bone metastasis was used via direct implantation of NSCLC cells (LLC and A549) into tibia of mouse. Mice were subsequently randomized into tumor-bearing only (control), treadmill running before and after implantation of NSCLC cells into tibia (exercise preconditioning), and treadmill running after implantation of NSCLC cells into tibia (treadmill running) groups. For exercise preconditioning group, mice were subjected to 4 weeks of treadmill exercise before implantation of NSCLC cells into tibia of mouse. Four weeks after implantation, mice were sacrificed.

b. Representative immunofluorescence staining images of GFP (green), CCND3 (red), and DAPI (blue) in the tibia of mice with bone metastases and quantification of the GFP and CCND3 positive cells. The dashed line indicates the boundary between bone and tumor. Scale bars: 50 μm. n=6, one-way analysis of variance with Turkey's multiple comparisons test.

**Supplemental Figure 7**

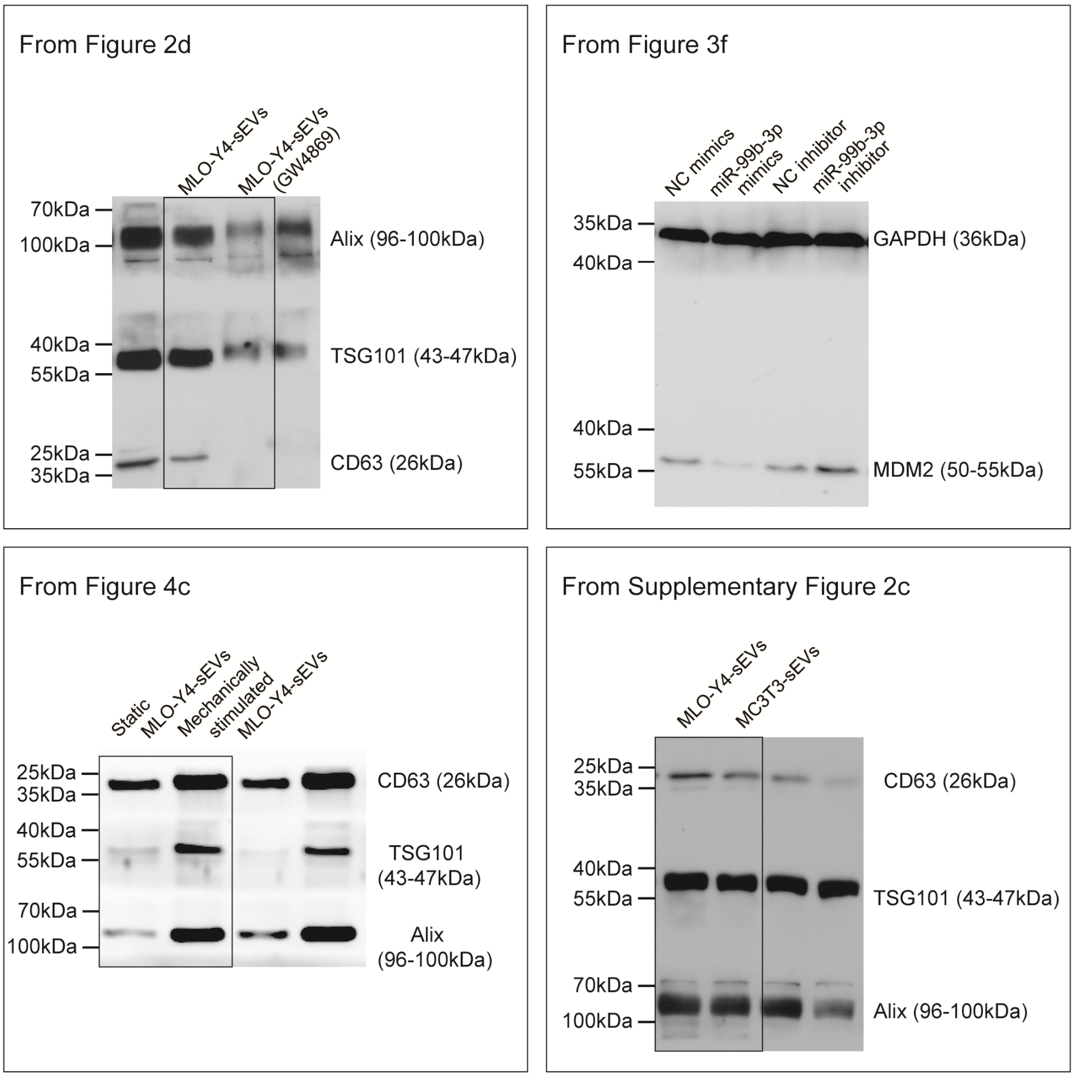


**Supplemental Figure 7. Uncropped images of western blots.**
